# Olanzapine: a full and potent agonist at the hM4D(Gi) DREADD amenable to clinical translation of chemogenetics

**DOI:** 10.1101/477513

**Authors:** Mikail Weston, Teresa Kaserer, Jenna C Carpenter, Albert Snowball, Samuel Knauss, Gabriele Lignani, Stephanie Schorge, Dimitri M Kullmann, Andreas Lieb

**Author notes:** these Authors contributed equally (Mikail Weston, Teresa Kaserer).

## Abstract

Designer receptors exclusively activated by designer drugs (DREADDs) derived from muscarinic receptors are a powerful tool to test causality in basic neuroscience, but are also potentially amenable to clinical translation. A major obstacle is however that the widely-used agonist clozapine-N-oxide undergoes conversion to clozapine, which penetrates the blood-brain barrier but has an unfavorable side effect profile. Perlapine has been reported to activate DREADDs at nanomolar concentrations, but is not approved for use in humans by the Food and Drug Administration or European Medicines Agency, limiting its translational potential. Here we report that the atypical antipsychotic drug olanzapine, widely available in various formulations, is a full and potent agonist of the human muscarinic-receptor M4-based DREADD, facilitating clinical translation of chemogenetics to treat CNS diseases.

## Introduction

CNS diseases caused by abnormal circuit function represent a major burden to society. Although many respond to conventional small molecule treatment, some diseases such as intractable pain and refractory epilepsy account for a substantial unmet need. Drug-resistant focal epilepsy alone affects approximately 0.2% of the entire population (*1, 2*). Although surgical resection of the epileptogenic zone is effective, it is contraindicated in the overwhelming majority of patients because of high risks of permanent disability associated with brain tissue removal (*3*). Several gene therapies for refractory epilepsy, based on altering the balance of excitation and inhibition, have been validated in preclinical models (*4-8*). Chemogenetics using viral vector-mediated expression of inhibitory DREADDs is especially promising because the therapeutic effect can be titrated by adjusting the dose of the activating ligand (*9*).

A potential limitation to clinical translation of DREADD technology is that most studies to date have used clozapine-N-Oxide (CNO), the inactive metabolite of the atypical antipsychotic drug clozapine (CZP) (*10*), as the ligand. CNO is not approved for clinical use, and recent evidence shows that CNO is actively exported from the CNS and back-converted to CZP, which crosses the blood brain barrier and subsequently acts as the ligand activating the DREADD (*11, 12*). CZP has a relatively high EC_50_ (~57nM) at the muscarinic receptor M4-based Gi-coupled DREADD (hM4D(Gi)) (*10*). This is up to 5-fold higher than the EC_50_ or IC_50_ at receptors relevant to its action as an antipsychotic drug (*13*). In addition, patients on CZP can develop agranulocytosis, myoclonus and generalized seizures (*14–16*). Although recent studies highlight the ability of hM4D(Gi) expressed in epileptogenic zones to suppress focal seizures when activated (*8, 17, 18*), the pharmacological profile of CZP is far from optimal for clinical translation. Related antipsychotic drugs have been proposed as potential agonists (*10, 11*), and two other drugs activating DREADDs have recently been described: “Compound 21” (C21) and perlapine (PLP) (*19, 20*). Although perlapine has previously been used as a mild sedative anti-histamine drug in Japan, neither it nor C21 is approved for clinical use by the FDA or EMA. Identification of an FDA/EMA-approved drug for repurposing as a DREADD activator would facilitate clinical translation of DREADD technology to treat CNS diseases.

## Results

### hM4D(Gi)-dependent Kir3.1 and Kir3.2 activation

In order to measure Gi-coupled hM4D(Gi) activation, we established an electrophysiological screen based on measuring the potentiation of the inward-rectifying potassium current in a HEK cell line stably expressing Kir3.1 and Kir3.2 (*21*) (**Fig. 1a**). We verified the sensitivity of the system by estimating the EC_50_ of CZP as 61±19nM (mean ± SEM, n=6), close to the reported EC_50_ of 57nM (*10*)). We used CNO (1µM) as a positive control to define maximal activation of hM4D(Gi) (**Fig 1b**), and confirmed that CZP, PLP and C21 are full agonists (efficacy in comparison to 1µM CNO: CZP/CNO=1.14±0.06, n=6; PLP/CNO=1.17±0.16, n=9; C21/CNO=1.11±0.07). C21 showed a significant lower EC_50_ than CZP (CZP EC_50_=61±19nM; PLP EC_50_=40±10nM; C21 EC_50_=20±4nM; p<0.05 one-way ANOVA with Bonferroni post-hoc test). We performed a shape (3D) (*22, 23*) and 2-D similarity (2D) screen (*24*) in order to identify FDA/EMA-approved drugs with structural and electrochemical properties similar to those of C21 (**Fig. 2a**). Prioritized drugs with similarity indicated by the TanimotoCombo score for the 3D screen, and by similarity for the 2D based screen, are listed in **Fig. 2b** and **Supplementary Table 1**.

**Figure 1:**
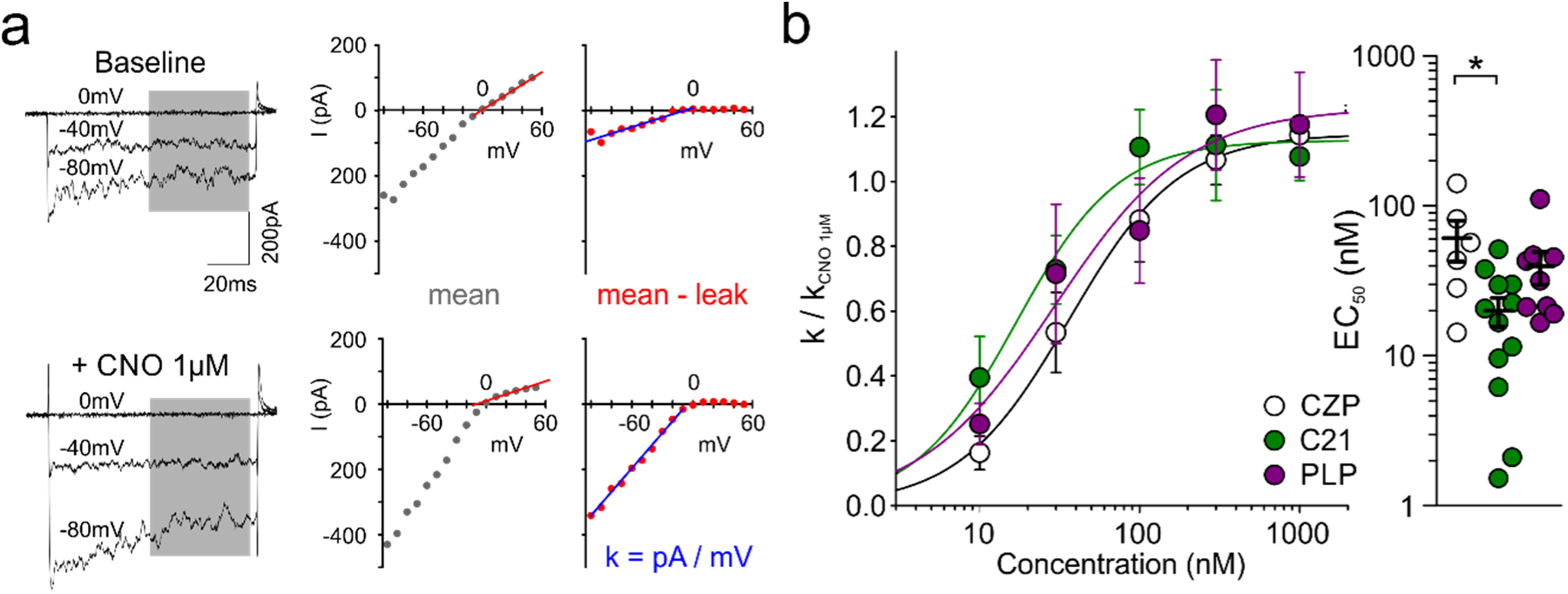
Electrophysiology-based screen of hM4D(Gi) activation. **a) Left:** Representative traces of Kir3.1 and Kir3.2 currents with (+CNO 1µM, lower) and without (baseline, upper) hM4D(Gi) agonist application. **Middle:** Mean current measured during the time indicated by the gray area in the left panel, plotted against holding voltage. The red line indicates the calculation of the membrane leak conductance, obtained from a linear fit between 0 and +50mV. **Right:** Leak-subtracted Kir3.1/Kir3.2-mediated currents, together with a linear fit to currents at negative potentials (blue). The slope of the current-voltage relationship (k) was used for subsequent analysis of hM4D(Gi) activation. **b) Left:** CZP, C21, and PLP act as full agonists of hM4D(Gi). All data are shown normalized to CNO (1µM) as a positive control, and fitted by a Hill equation. **Right:** EC_50_ of CZP, C21, and PLP (CZP: EC_50_=61±19nM, n=6; PLP: EC_50_=40±10nM, n=9; C21: EC_50_=20±4nM; * p<0.05 one-way ANOVA with Bonferroni post-hoc test).

**Figure 2:**
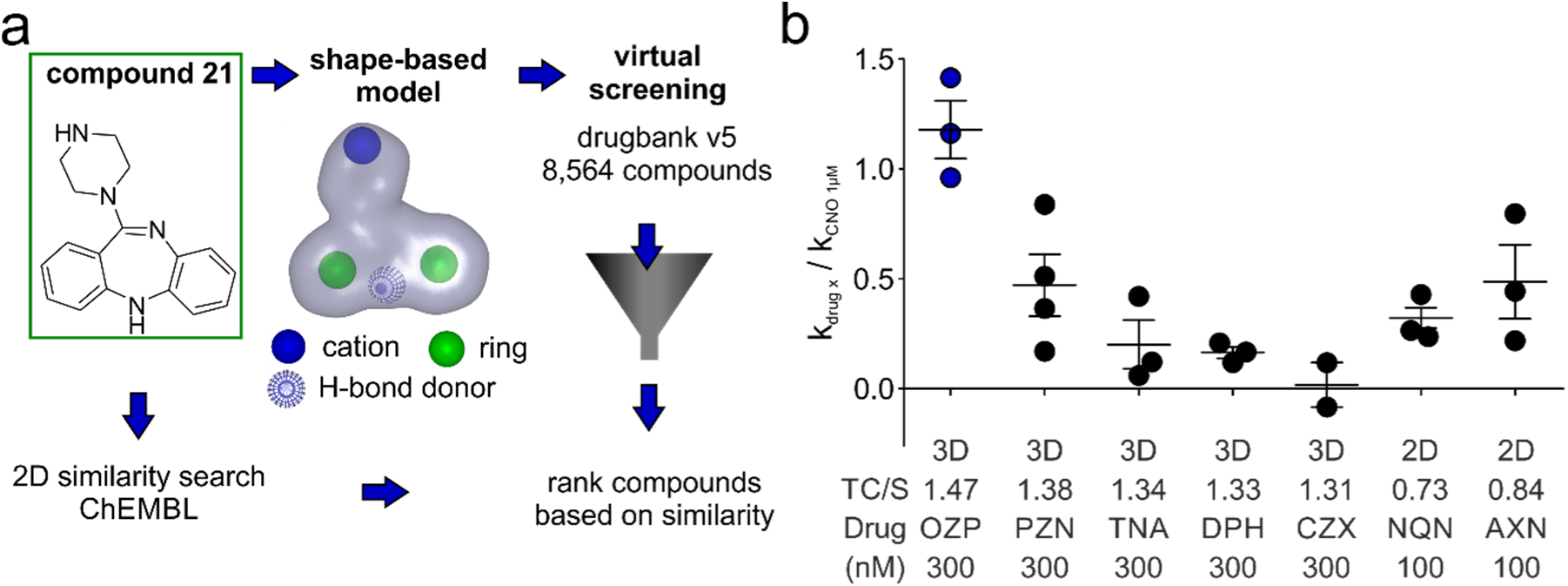
Shape- and 2D-similarity-based identification of novel hM4D(Gi) agonists. a) Overview of the 3D- and 2D-based virtual screen. C21 was used as the query compound for both the 2D-similarity search of the ChEMBL database and the generation of a 3D shape-based model, because it showed the lowest EC_50_ at hM4D(Gi) of known agonists. For detailed virtual screening results see **Supplementary Table 1**. b) hM4D(Gi)-dependent potentiation of Kir3.1/Kir3.2 mediated currents measured for selected hit compounds, normalized by 1µM CNO as positive control. The Tanimoto-Combo (TC) score (for 3D screen), similarity (S) index (for 2D screen), the screening method (3D or 2D), and the tested concentration is indicated. For detailed structures of selected drugs see **Supplementary Table2**.

### hM4D(Gi) activation with drugs identified by similarity screens

We tested olanzapine (OZP, 3D rank 1), promazine (PZN, 3D rank 2), triptilennamine (TNA, 3D rank 5), diphenhydramine (DPH, 3D rank 6), chlorprothixen (CZX, 3D rank 9), and amoxapine (AXN, 2D rank 2). We also tested the first, putative active metabolite of quetiapine (2D rank 6), norquetiapine (NQN) (*25*) (for chemical structures of all tested molecules – **Supplementary Table 2**). Of all drugs tested, only OZP was able to fully activate hM4D(Gi) at a concentration between 100-300nM, using 1µM CNO as control as above (OZP/CNO=1.18±0.13; n=3) (**Fig. 2b**). A full dose-response curve for OZP revealed an EC_50_ of 5±2nM (n=6), significantly lower than CZP (EC_50_=61±19nM, n=6; p=0.0128, Student’s t-test) (**Fig. 3a**).

**Figure 3:**
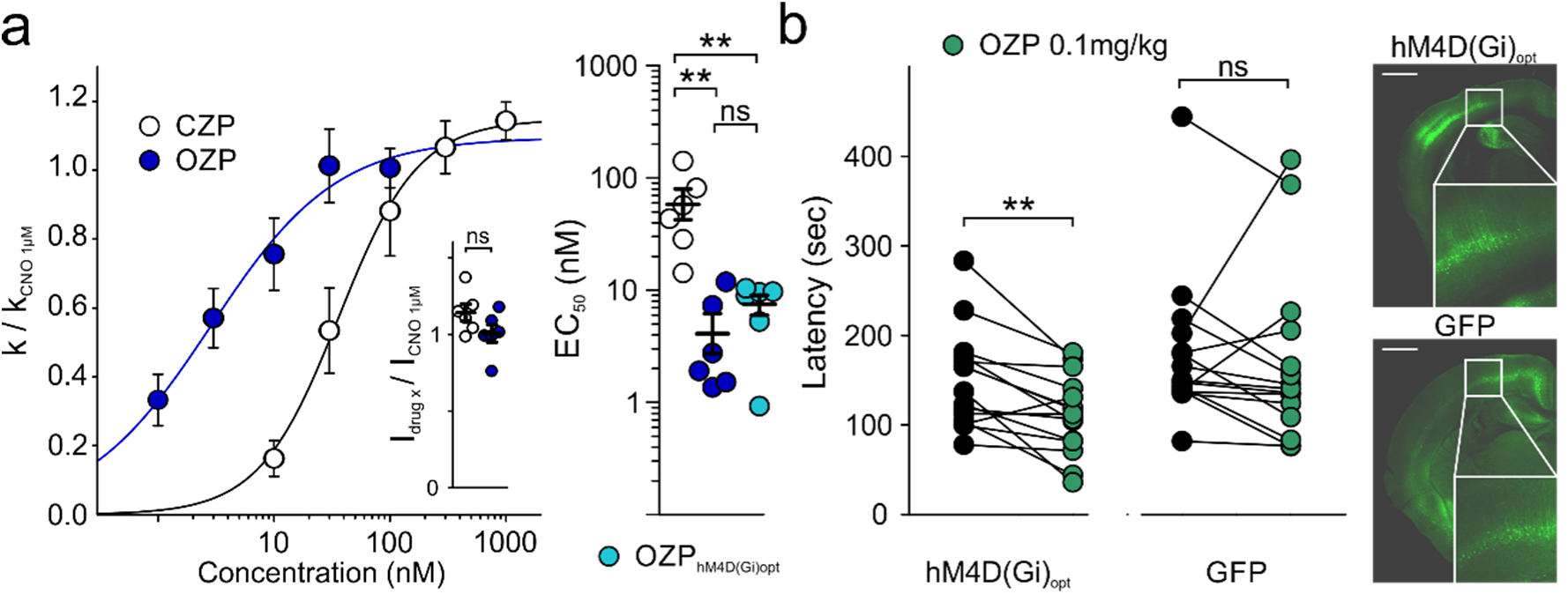
Olanzapine is a potent agonist at hM4D(Gi). **a) Left:** Dose-response curves for CZP and OZP at hM4D(Gi). The inset shows the efficacy of OZP (100nM) and CZP (1µM), normalized to 1µM CNO as a positive control (CZP/CNO=1.14±0.06, n=6; OZP/CNO=1.01±0.06; n=6; p=0.12, Student’s t-test). **Right:** EC_50_ for CZP at hM4D(Gi), and OZP at hM4D(Gi) and at the codon optimized hM4D(Gi)_opt_ (CZP: hM4D(Gi) EC_50_=61±19nM, n=6; OZP: hM4D(Gi) EC_50_=5±2nM, n=6, p<0.01 in comparison to CZP hM4D(Gi); OZP: hM4D(Gi)_opt_ EC_50_=7±2nM, n=6, p<0.01 in comparison to CZP hM4D(Gi); one-way ANOVA with Bonferroni post-hoc test). **b) Left:** Latency to fall (sec) of animals injected with either AAV2/8-hCamKII-hM4D(Gi)_opt_ (pre-OZP: 149±14sec, post-OZP: 111±11sec; **p=0.002, paired Student’s t-test), or AAV2/8-hCamKII-GFP (pre-OZP: 180±21sec, post-OZP: 169±25sec; p=0.568, paired Student’s t-test) pre- and post-injection of OZP (0.1mg/kg intraperitoneal). **Right:** Representative confocal fluorescence images of mouse brains injected with either AAV2/8-hCamKII-hM4D(Gi)_opt_ (DREADD), or AAV2/8-hCamKII-GFP (GFP) (scale bar 1mm).

### In vivo verification that olanzapine activates hM4D(Gi)

To test whether OZP is effective *in vivo* we redesigned a codon-optimized version of hM4D(Gi) linked via a viral self-cleaving 2A-peptide to GFP (hM4D(Gi)_opt_), and put it under control of a human CamKIIα promoter for selective expression in excitatory neurons (*26*). We verified that the EC_50_ of OZP at hM4D(Gi)_opt_ was similar to that at the original hM4D(Gi): OZP hM4D(Gi)_opt_: EC_50_=7±2nM, n=6; OZP hM4D(Gi): EC_50_=5±2nM, n=6). P0 mice were randomized for injection of either 2.5µl AAV2/8-hM4D(Gi)_opt_ or 2.5µL AAV2/8-CamKII-copGFP (control) into both lateral ventricles (**Fig. 3b right**). A third group of mice received no injection. After a period of training, their performance on the rotarod was then tested at ~P42, while blinded to the viral injection. All mice received an acclimatization session on the rotarod on the day of testing, followed by two test sessions, one before and one after ligand injection. We injected OZP at a dose of 0.1mg/kg intraperitoneally, approximately 10-fold lower than the dose reported to achieve a peak plasma concentration equivalent to that obtained in patients treated with a typical anti-psychotic dose (*27*). OZP significantly reduced the latency to fall from 149±14sec to 111±11 sec (n=15; p=0.002; paired Student’s t-test) in hM4D(Gi) injected animals. In contrast GFP-injected animals showed no significant difference pre- and post-OZP (control: 180±21, OZP 0.1mg/kg: 169±25; n=15; p=0.568; paired Student’s t-test) (**Fig. 3b**). OZP also had no effect in mice that had received no viral injection (control: 212±21, OZP 0.1mg/kg: 189±20; n=6; p=0.109; paired Student’s t-test).

## Discussion

Although recent papers highlight the potential of PLP or C21 as full and potent activators of the hM4D(Gi), they would require extensive screening to be approved for clinical use (*28*).PLP has been used clinically in Japan but was subsequently withdrawn from the market, calling for an alternative licensed drug that can be repurposed as an activator of hM4D(Gi) for clinical translation of DREADD technology. The present study shows that OZP (ranked first in the 3D-based *in silico* screen) is a full and potent activator of hM4D(Gi). Its EC_50_ in comparison to EC_50_ and IC_50_ at targets related to its clinical use as an antipsychotic, is lower than for CZP (*13*). OZP is a second-generation atypical antipsychotic which is approved by the FDA and EMA for treatment of schizophrenia and manic episodes in bipolar disorder. A common side effect of OZP at doses used in schizophrenia and bipolar disorder is a small degree of weight gain (*29*). OZP is a D2-receptor antagonist and its side effect profile therefore also includes akathisia, extrapyramidal symptoms, tardive dyskinesia, and neuroleptic malignant syndrome, although these are much less common than for first-generation antipsychotic drugs such as haloperidol and chlorpromazine. The *in vitro* EC_50_ of OZP at hM4D(Gi) is in the range of affinities reported for its native drug targets (*30*). The ability to affect performance on the rotarod with 0.1mg/kg OZP reported here is consistent with the principle of receptor reserve, whereby GPCR-mediated effects can be achieved with low doses of agonist (*31*). Given that CZP is typically only prescribed for treatment-resistant patients, due to its unfavorable side effect profile (*13*), CZP is much less suitable for repurposing as a DREADD activator. We therefore propose that OZP, which is widely available in oral, intramuscular and intravenous formulations, is suited for clinical translation of hM4D(Gi)-based chemogenetics to treat CNS diseases, including refractory epilepsy.

## Materials and Methods

### Voltage clamp recordings

A cell line stable expressing Kir3.1/3.2 (*32*) was cultured in DMEM GlutaMax^®^ (Gibco), supplemented with 10% fetal bovine serum (Gibco), Penicillin/Streptomycin (50 I.U./m, Gibco), and contained 500µg/ml Geneticin^®^ (Gibco) as a selection marker. Cells were transiently transfected with TurboFect^®^ transfection reagent (Thermo Fisher Scientific) with 3 µg of hM4D(Gi)-plasmid (Addgene, 45548) and 1 µg of CMV-GFP for cell identification. Standard whole-cell patch-clamp experiments were performed after 2-3 days as previously described (*26*). Briefly, borosilicate-glass electrodes were pulled (Sutter Instruments) and fire polished (Narishige), with a final resistance of 2-4.5 MΩ. The extracellular recording solution contained (in mM): KCl 140, CaCl_2_ 2.6, and MgCl_2_ 1.2, HEPES 10, adjusted to pH 7.4 with KOH. The intracellular recording solution contained (in mM): KCl 107, MgCl_2_ 1.2, CaCl_2_ 1, EGTA 10, HEPES 5, Mg-ATP 2, and NA_2_-GTP 0.3, adjusted with KOH to pH 7.2. Cells were voltage-clamped at a holding potential of 0 mV, and a 100 ms step depolarization from −100 mV to 50 mV was applied in 10 mV increments and a 30 sec inter-pulse interval. Whole cell currents were low-pass filtered at 2 kHz (Axopatch 1-D, Axon Instruments) and digitized at 10 kHz. The membrane leak conductance in each cell was estimated from a linear fit to currents measured between 0 and +50 mV. The inward rectifying conductance mediated by Kir3.1/3.2 was estimated from a linear fit to currents between −100 and 0 mV after subtracting the leak conductance (Fig. 1a). All recordings were performed at room temperature, and the different drugs were applied by a custom-built perfusion system. Clozapine-N-oxide (CNO, Generon, #HY17366), perlapine (PLP, Tocris, #5549), compound 21 (C21, HelloBio, #HB6124), clozapine (CZP, Cayman Chemicals, #12059), olanzapine (OZP, Santa Cruz Biotechnology, #sc-212469), promazine (PZN, Sigma Aldrich, #46674), triptilennamine (TNA, Santa Cruz Biotechnology, #sc-229608), diphenhydramine (DPH, Cerilliant, #D-015), chlorprothixen (CZX, Santa Cruz Biotechnology, #sc-211077), norquetiapine (NQN, BioVision, #2362), and amoxapine (AXN, LKT labs, #A5059) were dissolved in either DMSO or extracellular recording solution at a stock concentration of 1mM, and subsequently diluted to specified concentrations. CNO (1 µM) was routinely tested to estimate maximal activation of hM4D(Gi) in each cell, and the Kir3.1/3.2-mediated conductance activated by each agonist application was therefore related to that evoked by 1 µM CNO.

### Molecular biology

The hM4D(Gi)-plasmid was purchased from Addgene (#45548). Standard molecular biology techniques were used to clone dscGFP-T2A into an AAV2 transfer plasmid. The codon-optimized version of the HA-hM4D(Gi) (GeneOptimizer^®^, GeneArt^®^, Thermo Fisher Scientific) was linked to dscGFP via a viral 2A peptide. For in the *in vivo* experiments the CMV promoter was replaced with a CamKIIα-promoter to allow expression in excitatory neurons (*26*), the antibiotic resistance was changed from ampicillin to kanamycin, and a restriction site after the 2A peptide was removed.

### *In silico* screening

One low energy conformation of C21 calculated with Omega 2.3.2 (*33, 34*) was used as the query for the generation of the shape-based model. The default model was modified and the final model only contained the color features shown in Fig. 2a. A maximum number of 200 conformers were generated for Drugbank version 5.0.7 (*32*) with Omega 2.3.2 (*33, 34*). The default settings of vROCS 3.0.0 (*22, 23*) were used for screening and hits were ranked according to the TanimotoCombo score.

The ChEMBL (*24*) web service (https://www.ebi.ac.uk/chembl/; access date 2017/06/21) was employed to find FDA/EMA-approved drugs with similar 2D structure as C21.

### Viral injections

All animal procedures were performed in accordance with the [Author University] animal care committee’s regulations. Viral aliquots were prepared of AAV2/8-CamKIIα-GFP-T2A-hM4D(Gi)_opt_ or AAV2/8-CamKIIα-GFP (both titres >10^11^ GC/ml; Vectorbuilder) and coded by a researcher conducting neither surgical procedures nor behavioral analyses. P0 neonatal CL57BL/6 mice were anaesthetized with intraperitoneal ketamine 6mg/kg and midazolam 0.2mg/kg. A 10μl microinjection syringe fitted with a 32G angled needle (Hamilton) was filled with one or other virus. Mouse pups (n=30) were divided equally between viral types and manually injected with 2.5μl into each lateral ventricle, approximately 1mm lateral from the sagittal suture and halfway between lambda and bregma, to optimize widespread cerebral transduction. 6 pups received no injection. Pups’ paws were marked with green tattoo ink to allow differentiation between viral types, and after recovery they were returned to their home cage.

### Behavioral analysis

At P35 mice were trained on a mouse Rotarod (Ugo Basile). Mice were initially acclimatized for 10 minutes on the Rotarod turning at 5 revolutions per minute (RPM), replacing them each time they fell off, followed by acceleration over a 5-minute period from 5 to 40 RPM. The sequence was repeated 4 times. The latency to fall or to three consecutive cartwheels was recorded. Training was repeated daily until every mouse’s performance reached a plateau, taking approximately 2 weeks.

On the day of DREADD agonist testing the mice had a further acclimatization session (5 minutes at 5RPM followed by 4 accelerations) followed by a break of at least 30 minutes. They were then tested twice, with the same protocol as the acclimatization session, before and 20 minutes after intraperitoneal injection of OZP 0.1mg/kg. The latency to falling off or cartwheeling was recorded. Trial times were recorded using Excel (Microsoft), and statistical analyses performed with Graph Pad Prism 5.01, by a researcher blinded to viral treatment.

### Confocal Fluorescence

To establish the extent of viral transduction mice were anaesthetized with 150mg/kg pentobarbital (Boehringer Ingelheim) and transcardially perfused with 20ml heparinized (80mg/L) phosphate-buffered saline (PBS, Sigma-Aldrich) until the perfusate was clear, then switched to 4% paraformaldehyde (Tocris) (20ml). Brains were extracted and immersed in 4% PFA for a further 24 hours, before vibratome slicing (VT1000S Leica) at 50μm, mounting on slides with Vectashield mounting medium with DAPI (Vector Labs) and confocal fluorescence imaging (Zeiss LSM 710) to visualize GFP expression.

### Statistical Analysis

Statistical analysis was performed with Graph Pad Prism 5.01. Unpaired/paired Student’s t-test, or one-way ANOVA with Bonferroni post-hoc test was used as indicated. Data are shown as mean±sem, and the significant level was set to an α-error of p<0.05.

## Acknowledgments

We thank OpenEye for providing software free of charge. We are grateful to B. Roth for highlighting the potential of atypical antipsychotic drugs, and A. Tinker for the gift of the
Kir3.1/3.2 expressing cell line.

## Author contributions

A.L. and D.M.K designed all experiments and wrote the manuscript. A.L and S.K. performed and analyzed all *in vitro* experiments. T.K designed, performed, and analyzed all in silico experiments. M.W. and G.L. performed the vector injections, and M.W. performed and analyzed all *in vivo* experiments. J.C. and A.S. cloned the constructs. A.L., D.M.K., S.S., G.L., S.K., A.S., J.C., T.K., and M.W. revised the manuscript.

## Data and materials availability

All data are available upon request.

---

**Supplementary Table 1:** Hit list of the 3D- and 2D-based screens. Only FDA/EMA-approved molecules are listed (applicable for CHEMBL databank). The drug name, Tanimoto Combo (for 3D) or similarity (for 2D), as well as the screen database, and the individual identifiers (ID) are indicated. (Drugbank 5.0.7(*35*), CHEMBL(*24*), accessed 21.June 2017)

|  | rank | drug name | TanimotoCombo / similarity | database | ID |
| --- | --- | --- | --- | --- | --- |
| 3D | 1 | Olanzapine | 1.47 | drugbank | DB00334 |
| 3D | 2 | Promazine | 1.383 | drugbank | DB00420 |
| 3D | 3 | Chlorpromazine | 1.374 | drugbank | DB00477 |
| 3D | 4 | Clozapine | 1.363 | drugbank | DB00363 |
| 3D | 5 | Tripelennamine | 1.338 | drugbank | DB00792 |
| 3D | 6 | Diphenhydramine | 1.33 | drugbank | DB01075 |
| 3D | 7 | Chloropyramine | 1.32 | drugbank | DB08800 |
| 3D | 8 | Alimemazine | 1.312 | drugbank | DB01246 |
| 3D | 9 | Chlorprothixene | 1.305 | drugbank | DB01239 |
| 3D | 10 | Orphenadrine | 1.296 | drugbank | DB01173 |
| 2D | 1 | Clozapine | 88.9 | ChEMBL | ChEMBL42 |
| 2D | 2 | Amoxapine | 84.52 | ChEMBL | ChEMBL1113 |
| 2D | 3 | Perlapine | 80.34 | ChEMBL | ChEMBL340801 |
| 2D | 4 | Loxapine | 77.78 | ChEMBL | ChEMBL831 |
| 2D | 5 | Fluperlapine | 77.48 | ChEMBL | ChEMBL63756 |
| 2D | 6 | Quetiapine | 74.52 | ChEMBL | ChEMBL716 |
| 2D | 7 | Metiapine | 73.93 | ChEMBL | ChEMBL2106892 |
| 2D | 8 | Clothiapine | 72.85 | ChEMBL | ChEMBL304902 |

**Supplementary Table 2:**
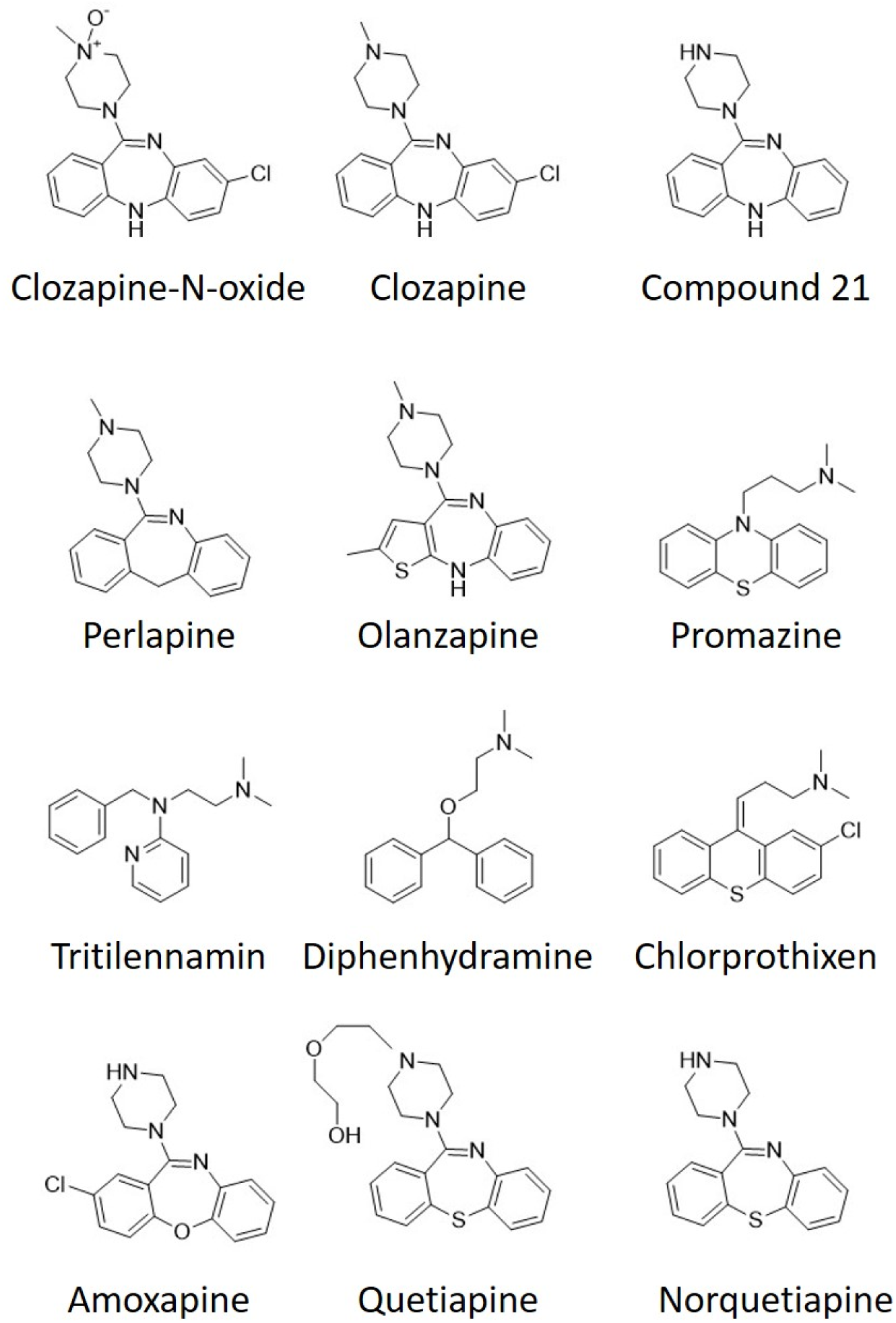
Structures of all tested molecules (note that also Norquetiapine is shown which has not been tested)

## References

1. p. Kwan, S. C. Schachter, M. J. Brodie, Drug-resistant epilepsy. The New England journal of medicine 365, 919–926 (2011).

2. F. Tang, A. M. S. Hartz, B. Bauer, Drug-Resistant Epilepsy: Multiple Hypotheses, Few Answers. Front Neurol 8, 301 (2017).

3. P. Ryvlin, J. H. Cross, S. Rheims, Epilepsy surgery in children and adults. The Lancet. Neurology 13, 1114–1126 (2014).

4. R. C. Wykes et al., Optogenetic and potassium channel gene therapy in a rodent model of focal neocortical epilepsy. Science translational medicine 4, 161ra152 (2012).

5. P. Kahane, A. Depaulis, Deep brain stimulation in epilepsy: what is next? Current opinion in neurology 23, 177–182 (2010).

6. S. M. Rothman, The therapeutic potential of focal cooling for neocortical epilepsy. Neurotherapeutics : the journal of the American Society for Experimental NeuroTherapeutics 6, 251–257 (2009).

7. J. D. Heiss, S. Walbridge, A. R. Asthagiri, R. R. Lonser, Image-guided convection-enhanced delivery of muscimol to the primate brain. J Neurosurg 112, 790–795 (2010).

8. D. Katzel, E. Nicholson, S. Schorge, M. C. Walker, D. M. Kullmann, Chemical-genetic attenuation of focal neocortical seizures. Nature communications 5, 3847 (2014).

9. A. Snowball, S. Schorge, Changing channels in pain and epilepsy: Exploiting ion channel gene therapy for disorders of neuronal hyperexcitability. FEBS letters 589, 1620–1634 (2015).

10. B. N. Armbruster, X. Li, M. H. Pausch, S. Herlitze, B. L. Roth, Evolving the lock to fit the key to create a family of G protein-coupled receptors potently activated by an inert ligand. Proceedings of the National Academy of Sciences of the United States of America 104, 5163–5168 (2007).

11. D. F. Manvich et al., The DREADD agonist clozapine N-oxide (CNO) is reverse-metabolized to clozapine and produces clozapine-like interoceptive stimulus effects in rats and mice. Scientific reports 8, 3840 (2018).

12. J. L. Gomez et al., Chemogenetics revealed: DREADD occupancy and activation via converted clozapine. Science 357, 503–507 (2017).

13. B. L. Roth, D. J. Sheffler, W. K. Kroeze, Magic shotguns versus magic bullets: selectively non-selective drugs for mood disorders and schizophrenia. Nat Rev Drug Discov 3, 353–359 (2004).

14. M. Sajatovic, H. Y. Meltzer, Clozapine-induced myoclonus and generalized seizures. Biol Psychiatry 39, 367–370 (1996).

15. S. Koch-Stoecker, Antipsychotic drugs and epilepsy: indications and treatment guidelines. Epilepsia 43 Suppl 2, 19–24 (2002).

16. C. J. Wenthur, C. W. Lindsley, Classics in chemical neuroscience: clozapine. ACS chemical neuroscience 4, 1018–1025 (2013).

17. E. Wicker, P. A. Forcelli, Chemogenetic silencing of the midline and intralaminar thalamus blocks amygdala-kindled seizures. Exp Neurol 283, 404–412 (2016).

18. R. Raedt et al., Chemogenetic silencing of excitatory hippocampal neurons prevents spontaneous seizrues in a mosue model for temporal lobe epilepsy. American Epilepsy Meeting, Abst. 1.080 (2016).

19. K. J. Thompson et al., DREADD Agonist 21 Is an Effective Agonist for Muscarinic-Based DREADDs in Vitro and in Vivo. ACS Pharmacology & Translational Science 1, 61–72 (2018).

20. X. Chen et al., The First Structure-activity Relationship Studies for Designer Receptors Exclusively Activated by Designer Drugs (DREADDs). ACS chemical neuroscience, (2015).

21. J. L. Leaney, G. Milligan, A. Tinker, The G protein alpha subunit has a key role in determining the specificity of coupling to, but not the activation of, G protein-gated inwardly rectifying K(+) channels. The Journal of biological chemistry 275, 921–929 (2000).

22. P. C. Hawkins, A. G. Skillman, A. Nicholls, Comparison of shape-matching and docking as virtual screening tools. J Med Chem 50, 74–82 (2007).

23. ROCS 3.0.0 OpenEye Scientific Software, Santa Fe, NM. http://www.eyesopen.com.

24. M. Davies et al., ChEMBL web services: streamlining access to drug discovery data and utilities. Nucleic Acids Res 43, W612–620 (2015).

25. N. H. Jensen et al., N-desalkylquetiapine, a potent norepinephrine reuptake inhibitor and partial 5-HT1A agonist, as a putative mediator of quetiapine’s antidepressant activity. Neuropsychopharmacology 33, 2303–2312 (2008).

26. A. Lieb et al., Biochemical autoregulatory gene therapy for focal epilepsy. Nature medicine 24, 1324–1329 (2018).

27. Y. E. Savoy et al., Differential effects of various typical and atypical antipsychotics on plasma glucose and insulin levels in the mouse: evidence for the involvement of sympathetic regulation. Schizophr Bull 36, 410–418 (2010).

28. M. J. Waring et al., An analysis of the attrition of drug candidates from four major pharmaceutical companies. Nat Rev Drug Discov 14, 475–486 (2015).

29. M. Huang et al., A randomized, 13-week study assessing the efficacy and metabolic effects of paliperidone palmitate injection and olanzapine in first-episode schizophrenia patients. Prog Neuropsychopharmacol Biol Psychiatry 81, 122–130 (2018).

30. F. P. Bymaster et al., Radioreceptor binding profile of the atypical antipsychotic olanzapine. Neuropsychopharmacology 14, 87–96 (1996).

31. B. L. Roth, DREADDs for Neuroscientists. Neuron 89, 683–694 (2016).

32. S. G. Brown, A. Thomas, L. V. Dekker, A. Tinker, J. L. Leaney, PKC-delta sensitizes Kir3.1/3.2 channels to changes in membrane phospholipid levels after M3 receptor activation in HEK-293 cells. American journal of physiology. Cell physiology 289, C543–556 (2005).

33. OMEGA 2.3.2: OpenEye Scientific Software, Santa Fe, NM. http://www.eyesopen.com. Hawkins, P.C.D.; Skillman, A.G.; Warren, G.L.; Ellingson, B.A.; Stahl, M.T.

34. P. C. Hawkins, A. G. Skillman, G. L. Warren, B. A. Ellingson, M. T. Stahl, Conformer generation with OMEGA: algorithm and validation using high quality structures from the Protein Databank and Cambridge Structural Database. J Chem Inf Model 50, 572–584 (2010).

35. D. S. Wishart et al., DrugBank 5.0: a major update to the DrugBank database for 2018. Nucleic Acids Res 46, D1074–D1082 (2018).

